# Circadian Rhythms of Whole-Body Metabolite Abundance in *Drosophila* are Largely Driven by Time of Feeding

**DOI:** 10.64898/2025.12.10.693378

**Authors:** Sumit Saurabh, Charlene Y. P. Guerrero, Madelyn R. Cusick, Amanda J. Samaras, Taylor M. Stephenson, Sebastian L. Kirkpatrick, Gregory J. Matthews, Daniel J. Cavanaugh

## Abstract

Organisms exhibit daily oscillations in metabolite abundance. These oscillations could arise from circadian clock control of metabolic pathways in peripheral tissues, or secondary to rhythmic food intake, which is primarily controlled by central circadian clocks in the brain. To determine the relative contribution of central and peripheral clocks and behavioral cycles to metabolic rhythms in the fruit fly, *Drosophila melanogaster*, we conducted large-scale metabolite profiling with fine temporal resolution across multiple days in control flies with intact molecular clocks and in flies in which we used the CRISPR/Cas9 gene editing system to specifically eliminate molecular circadian clock function in the fat body, a peripheral metabolic tissue, or the brain. As these latter flies lack feeding rhythms due to central circadian clock dysfunction, we also included an experimental cohort of flies lacking central brain clocks but subjected to time-restricted feeding (TRF) protocols to impose feeding rhythms. Single-nuclei RNA sequencing confirmed selective molecular clock elimination following fat body manipulations, which was associated with predicted alterations in clock gene expression and an attenuation of time-of-day differences in the abundance of fat body transcripts involved in key metabolic pathways. Interestingly, we identified few rhythmically expressed metabolites in flies that were allowed ad libitum food access, and rhythms of metabolite abundance were not drastically altered by tissue-specific molecular clock disruption. In contrast, we found that flies lacking brain clocks but raised on TRF exhibited a profound increase in cyclic metabolites. These findings suggest that whole-body metabolic rhythms in *Drosophila* are more strongly regulated by feeding cycles than by direct circadian clock control of metabolic pathways despite the presence of metabolic genes that exhibit local-clock dependent modulation of expression across the day.

## Introduction

As in other animals, the circadian clock in *Drosophila* consists of molecular components that form a ∼24-hr transcriptional-translational feedback loop (1). At its core, this involves the *Clock* (*Clk*) and *cycle* (*cyc*) genes, which encode for proteins that form a heterodimer and activate transcription of the negative regulators *timeless* (*tim*) and *period* (*per*). TIM and PER proteins accumulate in the cytoplasm and eventually translocate to the nucleus where they act as transcriptional repressors by inhibiting the function of CLK and CYC. Subsequent degradation of TIM and PER removes this repression, thereby initiating the next round of CLK/CYC-mediated transcription. A lack of genetic redundancy renders the *Drosophila* molecular clock dysfunctional following loss of any of these core clock molecules (2–5). In addition to the core feedback loop, a secondary loop involving rhythmic transcription of the *pdp1* and *vrille* (*vri*) genes generates cycles of *Clk* gene expression, thereby reinforcing circadian clock transcriptional rhythms (6).

The molecular clock functions in a dedicated group of central brain clock cells that generates rhythmic behavioral outputs (7). In addition, molecular clocks are found in cells in most peripheral tissues, where they impose time-of-day regulation on tissue-specific function (8). This is hypothesized to be achieved through clock-mediated oscillations of expression of hundreds of so-called clock-controlled genes, many of which are directly regulated by CLK/CYC binding to E-box elements in their promoters (9–11). The synchronization of circadian clocks across different tissues, which is likely facilitated by cross-tissue communication, serves to ensure optimal timing of behavioral and physiological outputs. For example, alignment between central and peripheral clocks coordinates feeding and activity rhythms with metabolic processes to maximize the efficiency of energy production, storage and use (12,13).

Consistent with circadian regulation of metabolism, previous studies have identified whole-body *Drosophila* metabolites whose levels are cyclically regulated across the day, and this cycling is altered in mutant flies with organism-wide loss of essential molecular circadian clock components (14–17). However, it is unknown whether *Drosophila* metabolic rhythms occur under the control of clocks in specific tissues. One important peripheral clock tissue in *Drosophila* is the fat body. Similar to mammalian liver and adipose tissue, the fat body serves functions related to metabolism and detoxification (18,19). The fat body is a primary location for intermediary metabolism, including protein synthesis and lipid and carbohydrate metabolism, and is a major storage site of energy reserves in the form of glycogen and triacylglycerides. In times of need, the fat body mediates the synthesis and mobilization of circulating metabolites such as trehalose, free fatty acids and diacylglycerides, which are used by target tissues to generate the energy that powers cellular functions (18,19).

Notably, fat body cells express components of the molecular circadian clock (20), and hundreds of genes have been found to exhibit circadian oscillations in the fat body (11,21). These include a number of genes involved in key metabolic processes performed by the fat body, such as the regulation of energy storage and release (22). Fat body gene expression rhythms appear to be under concurrent control of the local fat body clock and the central brain clock, indicative of the semi-hierarchical nature of the interactions between these tissues (21,23,24). At least some influence of the central brain clock on fat body gene expression rhythms could stem from entraining signals provided by systemic feeding cues, as feeding rhythms are strongly influenced by central brain clocks (21,24–26).

We hypothesized that metabolite abundance rhythms observed in flies arise from circadian regulation of energy metabolism pathways in the fat body. To test this, we conducted metabolite profiling in flies selectively lacking either central brain or fat body clocks, as well as in flies in which behavioral feeding rhythms were enforced through time-restricted feeding (TRF) paradigms. Through single-nuclei sequencing (snRNA-Seq) analysis, we confirm that the fat body circadian clock produces time-of-day-dependent differences in expression of a host of fat body genes that are enriched for those with metabolic functions. Nevertheless, we find that feeding time is the predominant factor regulating whole-body metabolite rhythms in *Drosophila*, with little apparent influence of fat body molecular clock cycling.

## Results

### CRISPR-Cas9-Mediated Molecular Clock Elimination

To assess the contribution of the fat body circadian clock to rhythms of whole-body metabolite abundance, we used the GAL4-UAS system (27) to drive expression of CRISPR-Cas9 constructs targeting the *per* gene (which we refer to as per^CRISPR^), thereby abrogating molecular clock function in a cell-specific manner (26,28). We first compared the effectiveness of two commonly used fat body driver lines, to-GAL4 and Lsp3.1-GAL4 (29,30), in inducing CRISPR-mediated elimination of PER protein expression in the fat body. In control flies, fat body cells were prominently labeled with PER antibody staining at zeitgeber time (ZT) 0, confirming molecular clock expression in these cells (Figure 1A) (20,26). Consistent with CRISPR-mediated excision of the *per* gene, PER staining was drastically reduced in fat body cells of flies in which either to-GAL4 or Lsp3.1-GAL4 was used to drive per^CRISPR^ expression (Figure 1A-C; Figure S1A-B). However, the effect was more complete with to-GAL4, which produced an ∼90% reduction in GFP-labeled fat body cells that co-expressed PER, compared to only a 67% reduction with Lsp3.1-GAL4 (Figure 1C). Induction of per^CRISPR^ with to-GAL4 or Lsp3.1-GAL4 did not appear to alter PER staining in molecular clock-containing oenocytes, nearby cells that also function in lipid metabolism (19,31,32) (Figure 1A-B; Figure S1A-B), in line with the cell-specific nature of the CRISPR ablation.

**Figure 1.**
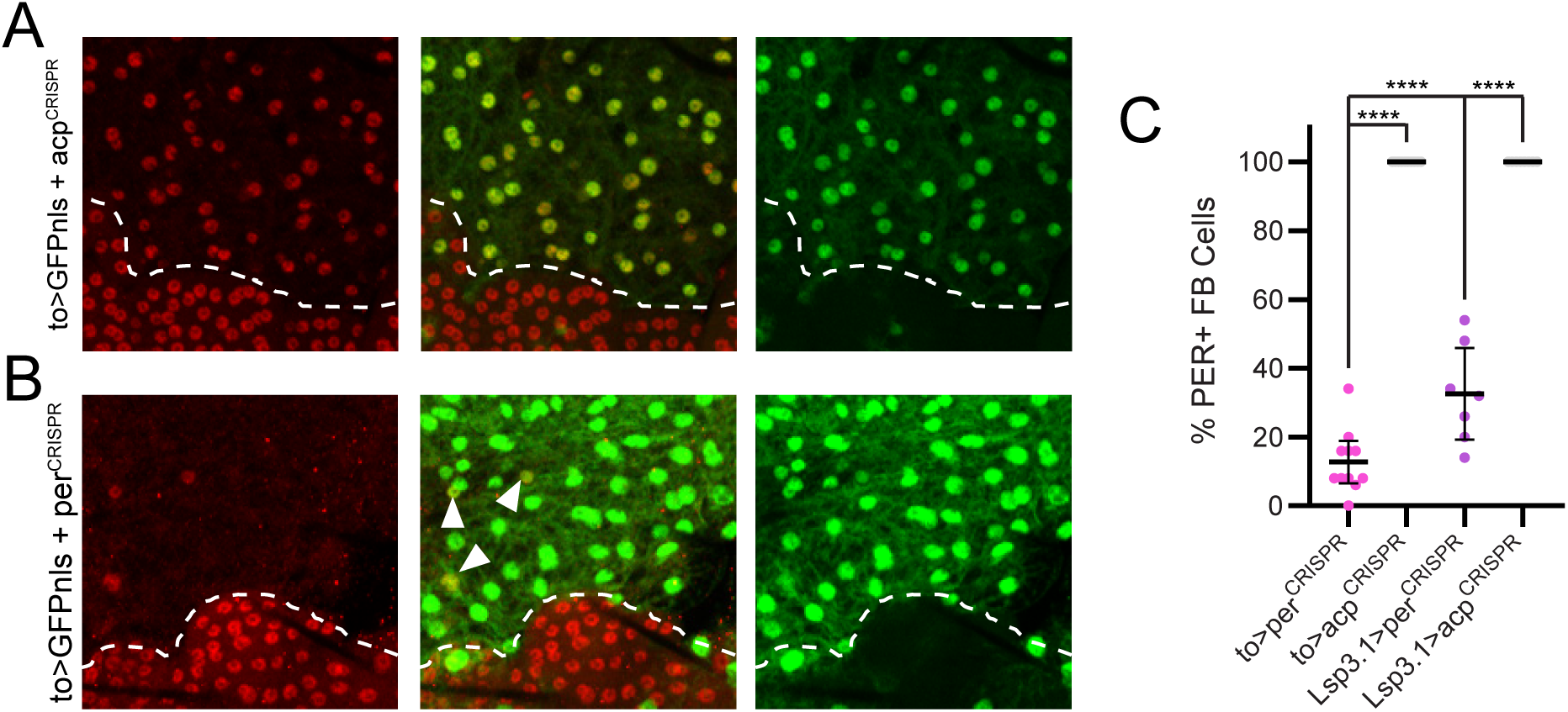
Selective elimination of fat body molecular clock. (A-B) Representative maximum projection confocal images of abdominal tissue (including fat body and oenocyte cells) stained for PER (red; left panels) and GFP (green; right panels) immunofluorescence. Center panels show merged images. (A) to>GFPnls + acp^CRISPR^ control sample shows co-expression of GFP and PER in fat body cells (above dashed line). Oenocytes (below dashed line) express PER but not GFP. (B) to>GFPnls + per^CRISPR^ experimental sample shows a loss of PER in GFP-expressing fat body cells (above dashed line), but retained PER expression in oenocytes (below dashed line). Arrowheads point to fat body cells that retained PER expression. (C) Quantification of the % of GFP-expressing fat body cells that also express PER. n = 7-11 samples per genotype. ****, p < 0.00001, Tukey’s multiple comparisons test.

In addition to testing for completeness of CRISPR effects, we also assessed specificity of the GAL4 lines for the fat body. Both to-GAL4 and Lsp3.1-GAL4 drove extensive expression of a nuclear GFP (GFPnls) reporter in adult abdominal fat body cells (Figure 1A-B; Figure S1A-B); however, they also showed additional expression outside the fat body. This extra-fat body expression was more extensive for to-GAL4 and included cells in the crop, cardia, hindgut, and rectal ampulla (Figure S1C). Notably, however, we found limited PER expression in these non-fat body to-GAL4-expressing cells (Figure S1D-G), indicating that the effect of to-GAL4-mediated clock disruption is relatively specific for the fat body. Thus, given the selective and nearly complete effect on PER elimination in the fat body, we decided to use to-GAL4 for subsequent experiments.

### Single-Nuclei RNA Sequencing (snRNA-Seq) Confirms Selective Fat Body Clock Disruption

These PER staining experiments confirm that per^CRISPR^ is effective in *per* gene disruption; however, the cellular circadian clock comprises multiple molecular components that interact to regulate cyclic transcription of many clock-controlled genes in addition to *per*. To assess the broader effects of fat body-induced per^CRISPR^ expression, we performed snRNA-Seq at two timepoints, circadian time (CT) 0 and CT12, that represent two distinct phases in the molecular clock oscillatory cycle: CT12 is a time of active CLOCK/CYCLE-mediated transcription whereas CT0 is a time of strong PER/TIM-mediated transcriptional repression. Our samples were taken from whole abdominal cuticle, which includes multiple distinct cell types, including fat body cells, oenocytes, and abdominal muscle cells (33). For these experiments, we compared transcriptomic profiles between to>per^CRISPR^ flies, which lack fat body molecular clocks, and to>Cas9 control flies, which omit specific guide RNAs necessary for CRISPR gene targeting. This enabled us to ascertain the direct effects of to-GAL4-mediated per^CRISPR^ expression in fat body cells, as well as potential indirect effects in other cell types not directly exposed to per^CRISPR^.

Our initial analysis identified 6 unique transcriptomic clusters at Leiden resolution of 0.2, and prominent among these were 3 large groups that we assigned to fat body, oenocytes and muscle cells based on known marker gene expression (33,34) (Figure S2A). The remaining smaller clusters are likely hemocytes, and digestive and reproductive tract cells from tissue remnants not fully removed during dissections (33). We found that these smaller clusters, as well as those we annotated as muscle cells, simultaneously expressed mRNAs of marker genes associated with multiple cell types (Figure S2B), perhaps due to ambient RNA or doublet contamination. Such cells have been recognized in previous single-nuclei datasets (34). We therefore focused our analysis on the fat body and oenocyte clusters, which more reliably expressed specific marker genes (Figure S2B-I).

To further investigate these two cell types, we performed dimensionality reduction using only nuclei that we annotated as either fat body or oenocytes based on initial clustering (Figure 2A). In this analysis, we found that fat body cells comprised multiple clusters (at Leiden resolution 0.5) that segregated by genotype, indicating that induction of per^CRISPR^ within these cells substantially altered their transcriptional landscape (Figure 2A-B). Interestingly, we also observed multiple oenocyte subclusters, with clear separation by genotype at CT12, indicative of non-cell-autonomous effects of molecular clock elimination within the fat body (Figure 2A-B).

**Figure 2.**
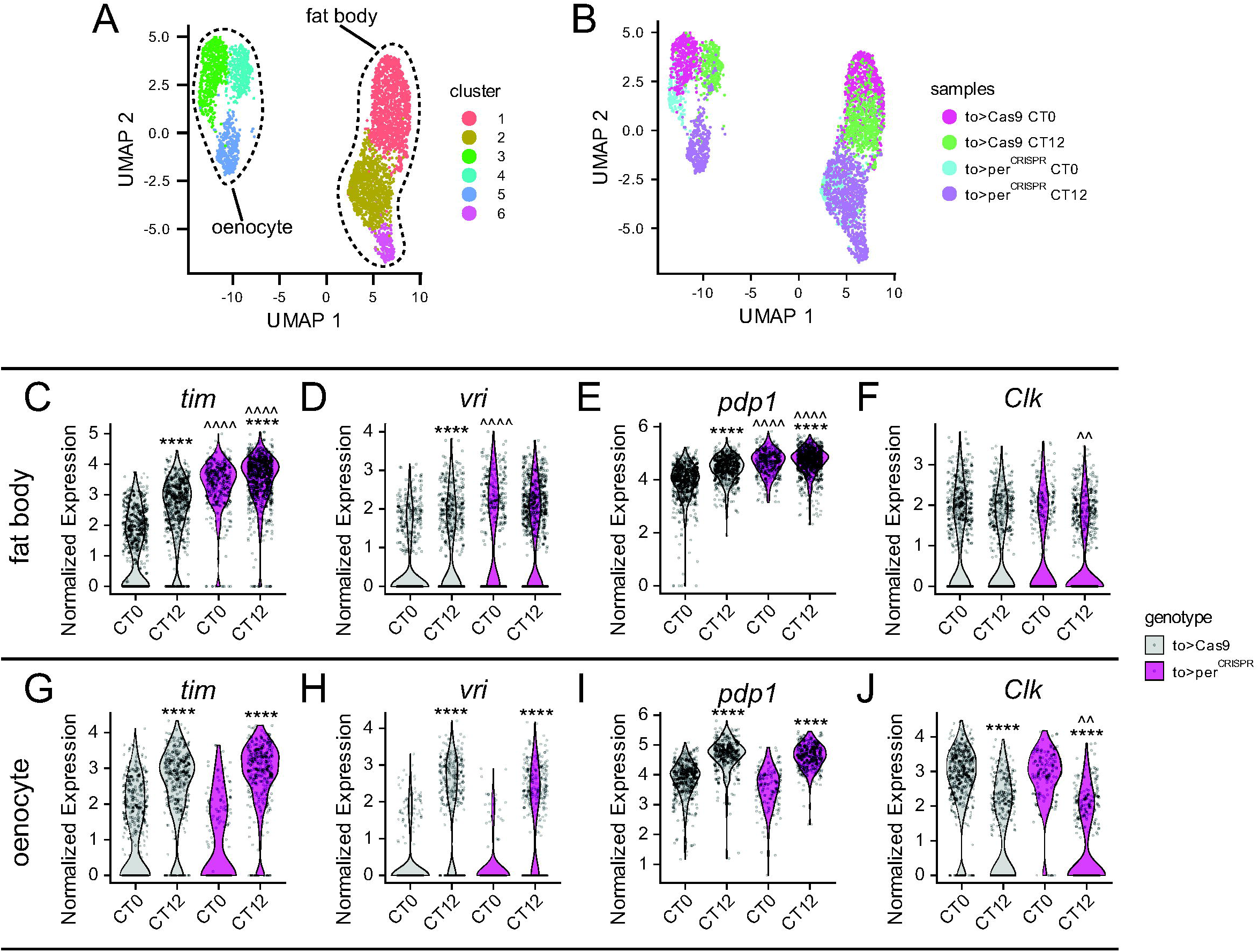
snRNA-Seq demonstrates dampening of fat body molecular clock cycling in to>per^CRISPR^ flies. (A-B) UMAP plots of nuclei annotated as fat body or oenocytes show 6 unique clusters at 0.5 Leiden resolution (A) that segregate by genotype and time point of samples (B). (C-F) Violin plots of normalized single nuclei transcript expression of core molecular clock components *tim* (C), *vri* (D), *pdp1* (E) and *Clk* (F) in fat body cells at CT0 and CT12 in control to>Cas9 (gray) and experimental to>per^CRISPR^ (magenta) flies. (G-J) Violin plots of core molecular clock transcript expression, as described for C-F, but in cells annotated as oenocytes. For C-J, each dot represents an individual nucleus. ****, *p* < 0.0001 compared to CT0 timepoint for the same genotype; ^^^^, *p* < 0.0001; ^^ *p* < 0.01 compared to to>Cas9 nuclei at the same timepoint, Memento differential expression analysis. Note the dampening of clock gene expression differences between CT0 and CT12 timepoints specifically in fat body cells of to>per^CRISPR^ flies.

Expression of per^CRISPR^ constructs in the fat body was associated with tissue-specific alteration of core molecular clock gene expression. We did not reliably detect significant levels of *per* expression in any sample at the timepoints and sequencing depth (∼60,000 reads/cell) that we used, precluding investigation of direct CRISPR effects on the *per* gene. Nevertheless, analysis of other core clock genes indicated a successful dampening of fat body molecular clock cycling. We performed differential expression analysis with Memento (35) to compare transcript levels of core clock genes within and between genotypes. In control to>Cas9 flies, we observed significantly lower expression of *tim*, *vri* and *pdp1* transcripts at CT0 compared to CT12 in cells annotated as fat body (Figure 2C-E), but *Clk* transcript levels were not significantly different between timepoints (Figure 2F). Consistent with a reduction of PER/TIM-mediated repression, abundance of *tim*, *vri* and *pdp1* transcripts was increased in fat body cells of to>per^CRISPR^ flies compared to controls, most significantly at CT0. This resulted in either a drastic dampening (for *tim* and *pdp1*) or elimination (for *vri*), of differential transcript abundance between CT0 and CT12 in to>per^CRISPR^ flies (Figure 2C-E). In contrast to the effect of per^CRISPR^ expression on fat body cells, expression levels and cycling of *tim*, *vri*, *pdp1* and *Clk* transcripts were largely unaffected in oenocytes (Figure 2G-J). While additional timepoints are necessary to confirm the presence or absence of oscillatory rhythms of expression, these data strongly support the effectiveness and specificity of per^CRISPR^ function in the fat body. They furthermore demonstrate coherent alteration of multiple core clock transcripts, consistent with a clock in which CLOCK/CYC is constitutively active.

### Autonomous Circadian Clock Regulation of Fat Body Metabolic Genes

A disrupted molecular clock should alter expression or cycling of clock-controlled genes in addition to the core clock genes we investigated above. We identified a total of 173 transcripts with differential abundance between CT0 and CT12 in fat body cells of control to>Cas9 flies (Figure 3A). Gene ontology analysis of these differentially-expressed fat body transcripts revealed an enrichment of those with metabolic functions, including carbohydrate, lipid and glycogen metabolism (Figure 3B). Notably, many of the differentially expressed genes that we identified have been previously implicated in fat body function. These include *Gart* (Figure 3C), which has been demonstrated to cycle in the fat body under control of the molecular circadian clock and to control triglyceride and glycogen levels, in addition to regulating circadian feeding rhythms (36). We also found differential abundance of transcripts of a number of lipid-regulating genes, including *brummer* (*bmm*), the major fat body lipase (37) and *Lsd-1*, which encodes for a lipid-droplet associated protein that regulates lipolysis (38) (Figure 3D-E).

**Figure 3.**
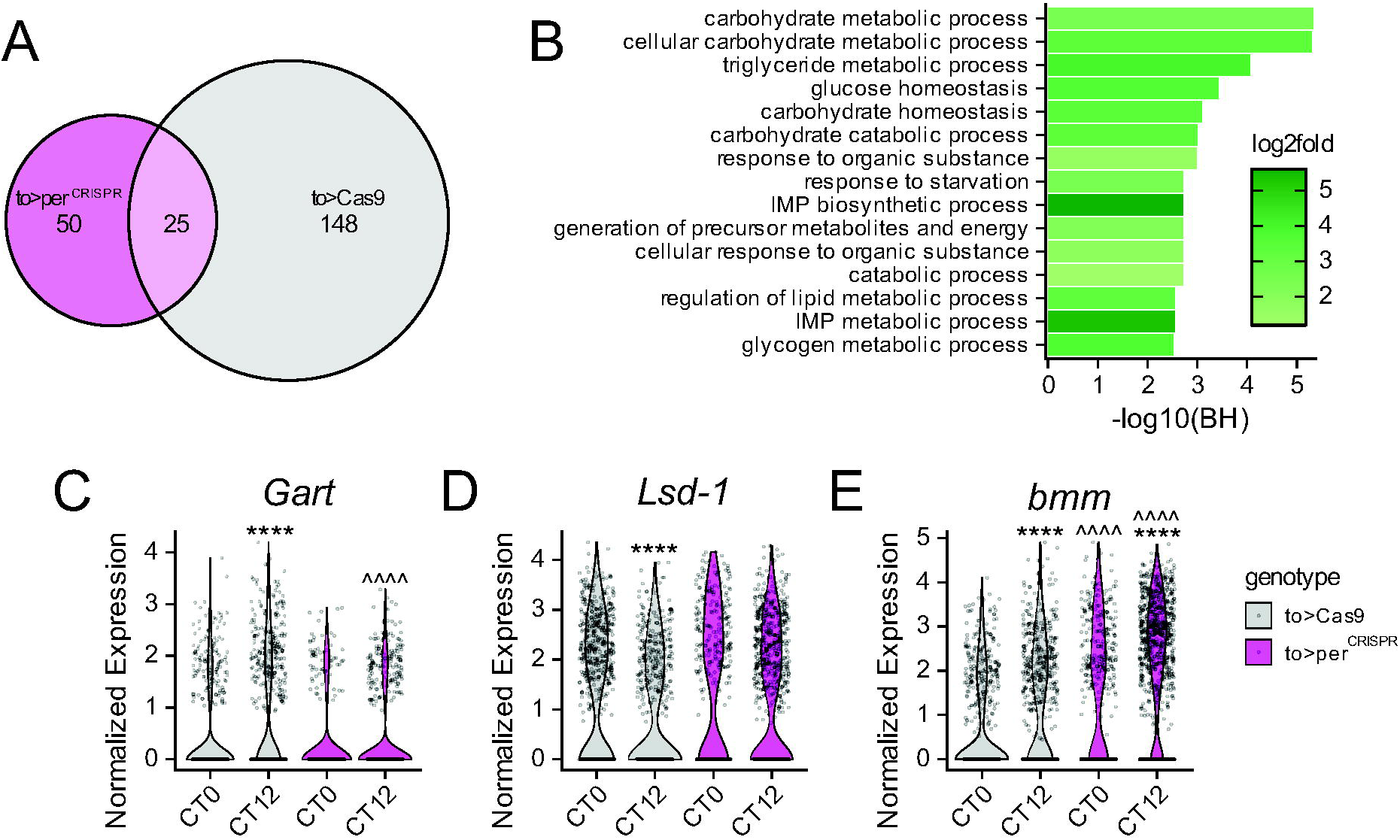
Time-of-day differences in fat body metabolic gene expression are reduced by fat body-specific molecular clock elimination. (A) Number of fat body transcripts with differential expression (at *p* < 0.05) between CT0 and CT12 based on Memento differential expression analysis of snRNA-Seq data. (B) Gene set enrichment analysis of differentially expressed fat body transcripts from to>Cas9 control flies showing the 15 most significantly enriched Gene Ontology biological process terms. (C-E) Violin plots show examples of metabolic genes with time-of day differences in expression in fat body nuclei of control to>Cas9 flies (gray), where differential expression is either eliminated (for *Gart* and *Lsd-1*; C-D) or dampened (for *bmm*; E) in to>per^CRISPR^ flies (magenta). For C-E, each dot represents an individual nucleus. ****, *p* < 0.0001 compared to CT0 timepoint for the same genotype; ^^^^, *p* < 0.0001 compared to to>Cas9 nuclei at the same timepoint, Memento differential expression analysis.

Reflecting a loss of molecular clock transcriptional control, we observed a very significant decrease in the number of differentially expressed transcripts in fat body cells of to>per^CRISPR^ flies (Figure 3A). Thus, the intrinsic fat-body clock is influential in dictating gene expression rhythms within the fat body, though some genes exhibit non-cell-autonomous cycling (35). This includes a loss of differential expression of both *Gart* and *Lsd-1* transcripts (Figure 3C-D), and a substantial reduction in the cycling amplitude of *bmm* (Figure 3E). These data demonstrate that the fat body molecular clock generates differential expression of metabolic genes, which could contribute to clock-controlled rhythms of metabolic function.

### Behavioral Effects of Elimination of Molecular Clock in Brain or Fat Body

Having established the molecular and genetic consequences of per^CRISPR^-mediated circadian clock abrogation in the fat body, we next sought to determine the behavioral effects of these manipulations. We used the DAM and FLIC systems to measure locomotor activity and feeding behavior of flies lacking either brain or fat body molecular clocks. Consistent with our previous analyses, we found that to-GAL4>per^CRISPR^ flies lacking fat body clocks exhibited normal free-running locomotor activity and feeding rhythm patterns under conditions of constant darkness, which was associated with a maintenance of rhythm strength as assessed by Lomb-Scargle periodogram (Figure 4A-D). In contrast, both activity and feeding rhythms were severely disrupted in Clk856>per^CRISPR^ flies, which lack central brain clocks (Figure 4A-D). Interestingly, however, Clk856>per^CRISPR^ flies showed some residual rhythmicity over the first ∼2-d of monitoring, with clear group-level activity and feeding peaks during the first and second subjective daytime periods (Figure 4A and C, insets) before completely losing rhythmicity. To our knowledge, such residual behavioral rhythms have not previously been reported in Clk856>per^CRISPR^ flies, which we suggest could arise either from a transient contribution from non-neuronal clock tissues, or to persisting effects of light exposure during entrainment.

**Figure 4.**
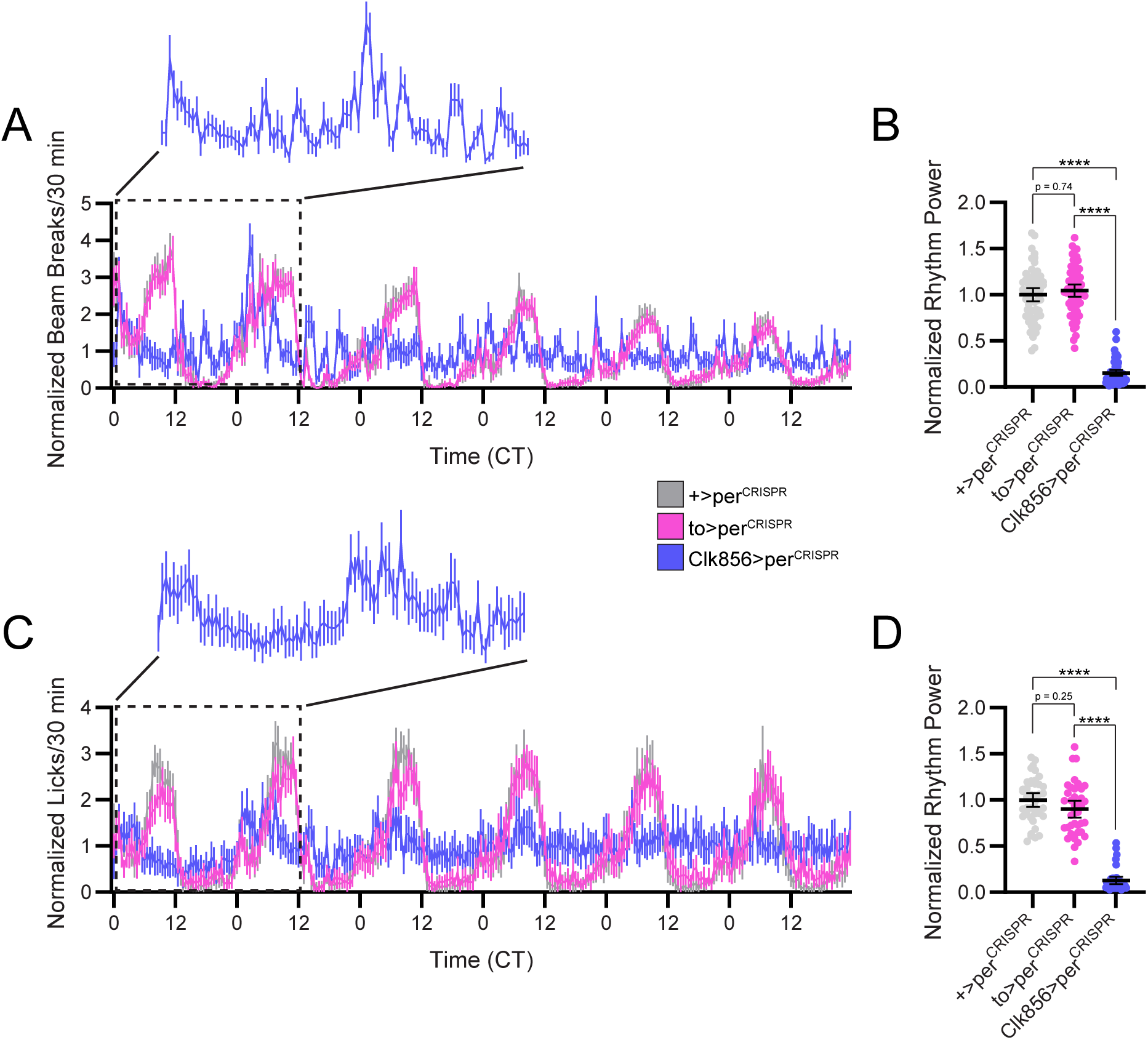
Locomotor activity and feeding rhythms are unaffected by loss of fat body molecular clock. (A) Mean locomotor activity (normalized DAM beam breaks/30 min ± 95% confidence interval) is plotted for 6 days in DD conditions for +>per^CRISPR^ (gray), to>per^CRISPR^ (magenta), and Clk856>per^CRISPR^ (blue) flies. Inset shows locomotor activity for the first 2 DD days (dashed rectangle) for Clk856>per^CRISPR^ flies, demonstrating transient rhythmicity during this time. (B) Normalized Lomb-Scargle locomotor activity rhythm power shows a selective reduction in Clk856>per^CRISPR^ flies. (C) Mean feeding (normalized FLIC licks/30 min ± 95% confidence interval) is plotted for 6 days in DD conditions for +>per^CRISPR^ (gray), to>per^CRISPR^ (magenta), and Clk856>per^CRISPR^ (blue) flies. Inset shows feeding behavior for the first 2 DD days (dashed rectangle) for Clk856>per^CRISPR^ flies, demonstrating transient rhythmicity during this time. (D) Normalized Lomb-Scargle feeding rhythm power shows a selective reduction in Clk856>per^CRISPR^ flies. For B and D, dots represent individual animals; lines are means ± 95% confidence intervals. ****, p < 0.0001, Dunnett’s T3 multiple comparisons test. n = 38-62 per genotype.

### Whole-Body Metabolite Abundance Rhythms are Largely Driven by Time of Feeding

Finally, we conducted metabolomics analysis to determine the relative contribution of molecular clocks and behavioral feeding rhythms to cycles of metabolite abundance. We assessed metabolite abundance rhythms in four groups of flies that differed in their molecular clock function and/or feeding behavior. These included +>per^CRISPR^ control flies with intact molecular clocks and to>per^CRISPR^ and Clk856>per^CRISPR^ flies, which lack fat body and central brain clock function, respectively. In addition to these three groups, which were allowed ad libitum food access, we also collected samples from Clk856>per^CRISPR^ flies subjected to time-restricted feeding, with food access limited to the 12-hour subjective day period. We collected whole-body samples every two hours over two days in DD conditions (with 2 experimental replicates per treatment group and timepoint) and then performed GC-TOF-MS metabolomics analysis to quantify metabolite abundance. Overall, we reliably detected 164 known and 412 unknown metabolites across our 4 treatment groups.

We then used the RAIN algorithm (39) to identify metabolites with circadian abundance profiles, restricting analysis to the 164 known metabolites. For each metabolite, RAIN tests for rhythmicity with multiple possible phases and waveforms and reports an adjusted *p*-value (adj-p) that corrects for the total possible combinations of phase and waveform. We then calculated Benjamini-Hochberg q (BH-q) values from these adj-p value to correct for the total number of metabolites tested per treatment group. Surprisingly, we did not detect robust metabolite abundance rhythms in control +>per^CRISPR^ flies. Although 10 metabolites were determined to be cyclically abundant in these flies with adj-p < 0.05, none of these retained significance after Benjamini-Hochberg correction (BH-q values > 0.05) (Figure 5A; Table S1). Similar results were observed in to>per^CRISPR^ and Clk>per^CRISPR^ flies under ad libitum feeding conditions, with BH-q values above 0.05 for all metabolites (Figure 5A; Table S1).

**Figure 5.**
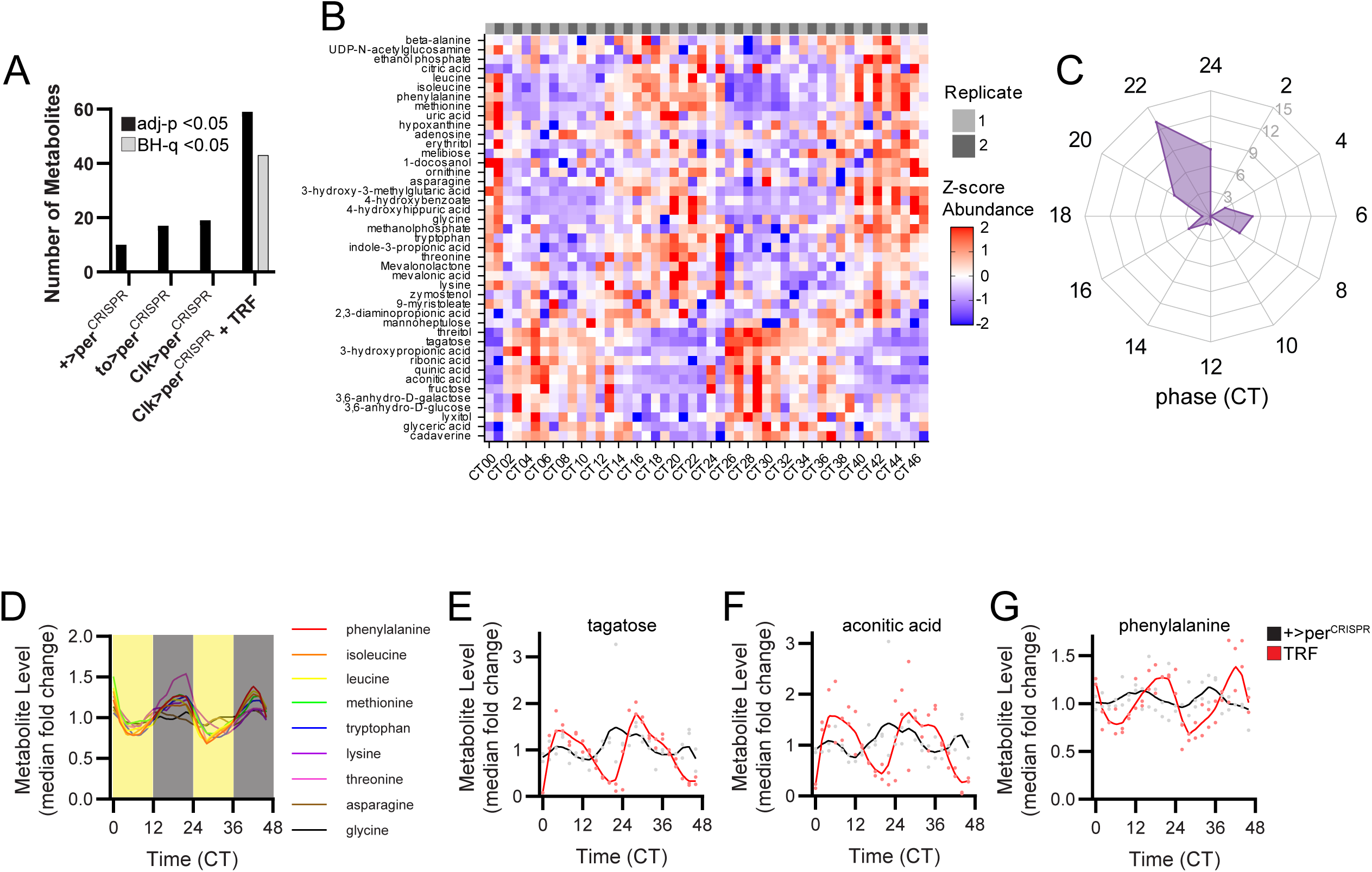
TRF increases the number of metabolites with circadian abundance rhythms. (A) Comparison of the number of metabolites found by RAIN analysis to have 24-hr rhythms with an adjusted-p value cutoff of 0.05 (black bars) or a BH-q false discovery rate cutoff of 0.05 (gray bars). (B) Heat map showing z-scored median fold change abundance of metabolites with RAIN BH-q values < 0.05 in Clk>per^CRISPR^ + TRF flies. Each timepoint consists of 2 experimental replicates. (C) Radial plot showing the number of rhythmic metabolites (radial axis) according to phase of peak abundance (angular axis), as determined by RAIN analysis, in Clk>per^CRISPR^ + TRF flies. Metabolites tended to peak around CT6 or CT22. (D) Abundance levels (LOWESS-filtered median fold change data) every 2 hours over 2 DD days of amino acids found to cycle (with BH-q < 0.05) in Clk>per^CRISPR^ + TRF flies. Yellow and gray shading represent feeding and fasting times, respectively. Amino acids showed a concerted peak towards the end of the fasting period. (E-G) Abundance levels of tagatose (E), aconitic acid (F) and phenylalanine (G) in +>perCRISPR (black) and Clk>per^CRISPR^ + TRF (red) flies. Dots represent median fold change values for individual experimental replicates and lines are LOWESS filtered data. These metabolites had 24-hr rhythms in control flies (adjusted p-value < 0.05, RAIN), but with lower amplitude and distinct peak phase compared to flies exposed to TRF.

In contrast, we found 43 metabolites with 24-hr abundance rhythms (BH-q < 0.05) in Clk>per^CRISPR^ flies subjected to TRF (Figure 5A-B; Table S1), indicating a strong impact of time of feeding on metabolite abundance. Interestingly, we observed that rhythmic metabolites tended to peak at one of two times of day: one group showed peak levels in the middle of the feeding period (CT4-8), and a second, larger group peaked towards the end of the fasting period (CT20-24) (Figure 5B-C). The first group included several sugars, which could directly result from consumption of the carbohydrate-rich food. We found a large group of amino acids among later-peaking group (Figure 5D), possibly indicating that protein metabolism is temporally delayed from time of food intake.

Overall, the substantial increase in the prevalence and magnitude of cycling in TRF flies demonstrates the predominant role of feeding time in dictating metabolic rhythms. Of note, despite the lack of robust cyclers in control +>per^CRISPR^ flies, we found weak evidence of cycling (adj-p < 0.05; BH-q > 0.05) of several of the metabolites identified as strongly circadian in TRF flies, although with reduced amplitudes and distinct phases of peak abundance in the ad libitum fed flies (Figure 5E-G).

## Discussion

Circadian clocks are thought to confer daily oscillations in tissue-specific functions. Here, we combined transcriptomic and metabolomic analyses with cell-specific molecular clock abrogation to test the hypothesis that local fat body clocks regulate organism-wide metabolic rhythms. However, despite identifying a large number of fat body metabolic genes with local clock-dependent expression patterns, we observed few metabolites that were rhythmically present in flies allowed ad libitum food access. In contrast, nearly 1/3 of identified metabolites exhibited rhythmic abundance patterns in TRF flies subjected to enforced feeding-fasting regimens, demonstrating that time of feeding, rather than oscillations in intrinsic metabolic processes, primarily dictate whole-body metabolite levels across the day. These findings are consistent with those observed in humans and mice, for which many circulating metabolites show rhythmic patterns that track to behavioral and feeding cycles (40,41), indicating that food intake strongly impacts metabolite rhythms across species.

Cyclic metabolites in TRF flies tended to peak either around CT6, in the middle of the feeding period, or around CT22, towards the end of the fasting period. Notably, metabolites that peaked coinciding with food availability included a number of sugars, which is consistent with results reported by Rhoades et al. (15). Despite this, we did not detect cycling of trehalose, the principal circulating sugar in the fly (18), nor of glycolytic intermediates, suggesting that the daily oscillations in sugar abundance do not translate to rhythmicity in energy metabolic processes. In addition to these early-peaking metabolites, we also observed that amino acids tend to accumulate during the enforced fasting period, which is consistent with the dark-phase selective upregulation of amino acids in flies exposed to 24-hr light and temperature cycles (14). Amino acids have also been found to exhibit prominent circadian cycling in mammalian metabolomics studies (42). Interestingly, most of the cyclical amino acids that we detected are essential, again suggesting an important contribution of time of feeding.

If, as we suggest, rhythmic feeding imposes metabolite rhythms, one question is why we don’t observe more cycling metabolites in control flies, which display prominent feeding rhythms (Figure 4C-D). We note that control flies do appear to have weak rhythms of several metabolites observed to cycle in TRF flies (Figure 5E-G). Thus, endogenous feeding rhythms could suffice to produce rhythmic patterns of metabolite abundance, but these are not as strong as in restricted feeding conditions. The much stronger rhythms observed in TRF flies could occur for multiple reasons. First, TRF synchronizes feeding time across the individual flies that are combined into a single sample, which would be expected to reduce variability due to inter-animal differences. Second, TRF institutes a prolonged and complete fasting period throughout the subjective night, which could be important for producing robust metabolic cycling. The relatively greater influence of TRF compared to endogenous feeding rhythms on molecular cycling has been observed in other contexts. For example, in mice, natural feeding rhythms do not entrain peripheral clock gene expression to the same extent that TRF does (43).

Our failure to identify more robustly-cycling metabolites in control flies was unexpected, as several previous metabolomics studies have identified metabolites that exhibit circadian abundance patterns in *Drosophila* (14–17,44–46). However, relatively few specific metabolites have been shown to cycle in common across studies, perhaps due in part to the fact that the different studies used a variety of samples, lighting regimes, metabolite detection methods, and statistical tests of rhythmicity. In particular, the presence of entraining environmental cues, which were present in the majority of previous studies, could serve to strengthen internal metabolic cycles and synchronize metabolite levels across the multiple flies in a manner similar to that produced by TRF in the current study. In fact, several studies have reported that metabolite cycling in *Drosophila* is blunted or even eliminated under constant environmental conditions (17,47,48).

It is also possible that our whole-body metabolomics analysis is obscuring tissue-specific metabolic rhythms, as have been identified in mice (49). While it is technically difficult to isolate specific organs for metabolite profiling in flies, Amatobi et al. recently profiled lipid species (many of which would be expected to be produced in the fat body) circulating in fly hemolymph, and found that a large number of glycerolipids and phospholipids had circadian abundance patterns under LD conditions (47). These rhythms persisted but with reduced amplitude in DD. Our use of gas chromatography for metabolite extraction precluded identification of these lipid species, so we are unable to determine whether they cycle under our experimental conditions. In the future, it would therefore be of interest to conduct targeted lipidomics analysis on either isolated fat body or hemolymph to see if loss of the intrinsic fat body clock alters glycerolipid expression or cycling. We note, however, that hemolymph lipid abundance rhythms were severely blunted when flies were fed standard as compared to sucrose-only food (47). This could indicate that under nutritionally replete conditions, intrinsic lipid metabolism is less effective in dictating circulating lipid levels. Thus, in addition to restricting metabolomics analysis to individual tissues, it may be necessary to control dietary conditions to uncover circadian metabolic control.

In addition to metabolomic analysis, we conducted snRNA-seq to identify genes whose expression is regulated by the fat body clock. Notably, we found that the transcripts of many fat body genes, including an abundance of genes with metabolic function, exhibit differential expression between CT0 and CT12, and this was considerably altered by fat body-specific molecular clock ablation. This list of differentially-expressed transcripts showed a >7-fold enrichment for those previously determined to be rhythmically expressed in the fat body based on bulk RNA-sequencing analysis (11). This demonstrates the fidelity of our snRNA-seq approach in identifying circadianly-regulated genes, even with very limited temporal resolution. However, more timepoints are required to confirm that the differential expression we observe between CT0 and CT12 reflects de facto circadian control.

Surprisingly, this differential gene expression does not translate into prominent rhythms in metabolite levels. It is possible that gene expression cycles occur in anticipation of feeding, such that gene upregulation at specific times of day enhances the ability to efficiently metabolize incoming nutrients. Such an organization could serve to maintain consistent energy reserves across feeding and fasting times rather than to produce drastic changes in metabolite abundance throughout the day, as has been suggested for liver-clock-dependent maintenance of glucose homeostasis in mice (50). Interestingly, we found that lipids and related compounds represented 8/17 metabolites with adj-p < 0.05 but BH-q > 0.05 in to>per^CRISPR^ flies, and such compounds were largely arrhythmic in other groups (Table S1), potentially indicating that the fat body molecular clock prevents fluctuations in lipid abundance across the day.

Together, our results demonstrate the potential for prominent circadian clock control over fat-body dependent metabolic pathways and show that this stems from local fat body clock regulation of gene expression. However, the substantial time-of-day differences in the fat body transcriptomic landscape produce substantially more modest changes in metabolite levels, at least at the whole-body level. It is therefore possible that in some contexts, peripheral circadian clocks function to anticipate behavioral cycles to buffer feeding-fasting-dependent fluctuations in nutrient availability.

## Materials and Methods

### Fly Stocks

Flies were raised in narrow polystyrene vials (Fisher Scientific AS516) in incubators held at 25°C and running on a 12:12 light-dark (LD) schedule. Flies were provided a cornmeal-molasses medium consisting of (per L food): 1L deionized water, 64.7 g yellow cornmeal, 27.1 g dry active granular yeast, 8.0 grams 80-100 mesh agar, 90 g unsulphured molasses and supplemented with 4.4 mL propionic acid and 2.03 g Tegosept to prevent contamination. Iso31 (isogenic w1118) (51) and to-GAL4 (FBti0202314) were provided by Amita Sehgal. Clk856-GAL4 (52) was provided by Orie Shafer. Lsp3.1-GAL4 (FBti0210031) was provided by Brigitte Dauwalder. UAS-GFPnls (FBti0012492) and UAS-Cas9.P2 (FBti0166500) were provided by the Bloomington Drosophila Stock Center. UAS-sgRNA-per^4x^ and UAS-sgRNA-acp^4x^ were provided by Mimi Shirasu-Hiza. All fly lines were outcrossed at least 7 times to the Iso31 background.

### Immunohistochemistry

Adult male flies were entrained to a 12:12 LD cycle at 25°C for ≥ 7 d, and dissections were performed at ZT0. Flies were CO_2_ anesthetized, submerged for ∼1 min in ethanol and rinsed briefly in phosphate-buffered saline with 0.1% Triton-X (PBST) before abdomen and digestive tract dissection in PBST. Dissecting forceps were used to tear a small opening in the caudal end of the abdomen, which served as an access point for removing internal tissues such as reproductive and digestive tracts. The caudal abdomen was then isolated, and the posterior-most genital segment was removed. An incision was then made along the ventral abdominal surface to remove the ventral cuticle.

Following dissection, whole-mount abdomens (including fat bodies and oenocytes) and digestive tracts (including crop, rectal ampulla, Malpighian tubules, fore-, mid- and hindgut) were processed for immunohistochemical analysis. Tissues were fixed in 4% formaldehyde for 15–35 min, washed 3 × 15 min in PBST, blocked for 1 hr in 5% normal donkey serum in PBST (NDST) and incubated for 24 hrs in primary antibodies diluted in 5% NDST. They were then washed 3 × 15 min in PBST and incubated ∼24 hrs in secondary antibodies diluted in 5% NDST. Finally, tissues were washed 3 × 15 min in PBST and mounted in Vectashield (Vector Laboratories). Primary antibodies were guinea pig anti-PER 1:1000 (UPR 1140; gift of A. Sehgal) and rabbit anti-GFP 1:1000 (Molecular Probes A-11122). Secondary antibodies were Cy3 donkey anti-guinea pig 1:1000 (Jackson 706-165-148) and FITC donkey anti-rabbit 1:1000 (Jackson 711-095-152).

### Confocal Imaging and Analysis

To facilitate identification of fat body cells, we recombined either to-GAL4 or Lsp3.1-GAL4 with UAS-GFPnls. These to-GAL4, UAS-GFPnls and Lsp3.1-GAL4, UAS-GFPnls flies were crossed with UAS-acp^CRISPR^ (which targets the non-circadian *acp98* gene that is expressed in male accessory gland and testes) or UAS-per^CRISPR^ constructs and dissected abdomens were processed for PER and GFP immunofluorescence as described above. Immunolabeled samples were visualized with an Olympus Fluoview FV1000 confocal microscope. 20x images were taken of 10-11 abdomens per genotype, with a Z-stack through the entire dorsal-ventral axis of each abdomen. Gain and offset were set to minimize saturation, and confocal capture settings were held constant across all samples. PER knockout in fat body cells was quantified using the ImageJ cell counter plug-in. A 10 μm maximum projection image was created, and 50 GFP+ fat body cells were randomly selected. These 50 cells were then assessed for the presence of PER staining above background levels, and we determined the overall percentage of GFP-expressing cells in each sample that also expressed PER. For statistical comparison of the percentage of PER-expressing GFP cells across genotypes, one-way ANOVA with Tukey’s multiple comparisons test was performed with GraphPad Prism 10 software.

### Single-Nuclei RNA Sequencing

#### Sample Preparation and Nuclei Isolation

Nuclei from abdominal tissues were sampled at two circadian timepoints: CT0 and CT12. 7 d old control (to-GAL4>UAS-Cas9.P2) and experimental (to-GAL4>UAS-Cas9.P2; UAS-gRNA-per) flies were raised under 12:12 LD cycles and transferred to DD conditions on the day of tissue collection. Samples consisting of 30 pooled abdomens per genotype and timepoint were dissected in ice-cold Scheider’s medium (SM) following ∼1 min incubation in ethanol. Following dissection, abdomens were flash frozen in liquid nitrogen and stored in a minimal volume of SM at -80°C until nuclei isolation.

Nuclei isolation was performed using protocols adapted from McLaughlin et al. (53). Frozen fly abdomens were quickly thawed on ice and resuspended in 950 μl of homogenization buffer (HB) consisting of: 250 mM sucrose, 10 mM Tris (pH 8.0), 25 mM KCl, 5 mM MgCl_2_, 0.1% Triton X-100, 1x protease inhibitor cocktail (Promega G6521), 0.1 mM DTT, 200 U/mL RNAsin Plus (Promega N2615), and 20 U/mL heat-activated DNAse I (Thermo Scientific EN0521), in nuclease free water. Resuspended abdomens were transferred to a prechilled (4°C) Wheaton dounce homogenizer (1 mL capacity; Fisher Scientific 06-434) on ice and allowed to settle at the bottom of the homogenizer. To free nuclei, the loose pestle was first used to shear the tissue with 20 half-turns while trapping the abdomens at the bottom of the dounce. This was followed by 20 strokes with the loose pestle and 40 strokes with the tight pestle. The homogenized tissue was then filtered through a SWiSH 30 micron mini cell strainer (Stellar Scientific TC70-SWM-30) and centrifuged for 10 min at 500 x g at 4°C in a fixed-angle table-top centrifuge. Following isolation, the Evercode Nuclei Fixation V3 kit (Parse Biosciences ECFN3300) was used to fix and store nuclei following manufacturer instructions. To quantify and ensure nuclei integrity, acridine orange/propidium iodide (AO/PI) staining was performed on a 10 μl aliquot of fixed nuclei and a combination of brightfield and fluorescence microscopy was carried out with a Zeiss AXIO Imager microscope.

#### Library Preparation and Sequencing

Single nuclei barcoding and library preparation with the Evercode WT Mini V3 kit (Parse Biosciences ECWT3100) was performed by the Northwestern University Center for Genetic Medicine NUseq Core facility according to manufacturer instructions. Libraries were subjected to paired-end 150 base pair sequencing with the NovaSeq X Plus platform.

#### Transcriptomics Data Analysis

FASTQ files were processed for barcode correction, read alignment, read deduplication, and transcript quantification using Parse Biosciences Trailmaker^TM^ pipeline module (v1.5.1). The reference genome was generated by adding GAL4 and Cas9.P2 (54) sequences to the *Drosophila melanogaster* genome (*Ensembl* BDGP6.32 release 109) (55) Further processing of transcripts was done in the Trailmaker^TM^ insights module. Nuclei with <1000 transcripts, >0.2% mitochondrial content, and >0.2 probability of being a doublet (determined by scDblFinder) were filtered out. Outliers in the distribution of the number of transcripts vs. number of genes were removed, as determined by fitting a spline regression model with default settings. Data normalization, principal component analysis (PCA), and data integration were performed using Seurat and Harmony. Initial clusters were identified with the Leiden method at a 0.4 resolution, and a Uniform Manifold Approximation and Projection (UMAP) embedding was calculated.

The resulting data were further processed in R (v4.5.1) with the Seurat package (v5.2.0) (56) Ambient RNA contamination was filtered out using automatic estimations from SoupX (57) with the original unfiltered data and the clustered data from Trailmaker^TM^ used as input. Nuclei were removed if their transcript count was >3 absolute deviations from the median transcript count, and genes were removed if they were expressed in <3 nuclei. After this filtering, data were processed with the standard Seurat workflow, using the following functions with default values unless otherwise stated. Raw counts were normalized with NormalizeData, which divides transcript counts from each nucleus by the total number of counts for that nucleus, multiples them by a scaling factor, and performs a natural log normalization of the result. Highly variable genes were determined with FindVariableFeatures and normalized transcript counts were scaled with ScaleData so that PCA could be performed using RunPCA, and the first 10 PCs were used for downstream processing. New clusters were determined with FindNeighbors and FindClusters, using the Leiden algorithm at a 0.2 resolution. Finally, non-linear dimensional reduction was performed with RunUMAP. A subset of the data including only oenocytes and fat body cells, as determined by marker gene expression, was further clustered with the same Seurat workflow, retaining the first 10 PCs and using the Leiden algorithm at a 0.5 resolution, and used for subsequent analyses.

Differential expression analysis between genotypes and timepoints within a cell type was conducted using Memento (v0.1.2) (35) in Python (v3.12.2). Data were analyzed using the binary_test_1d method with a capture rate of 0.035 and all other settings as default. The de_coeff was used as effect size and the de_pval was corrected across all transcripts using the Bonferroni method in R, which was reported as the adjusted *p*-value. We considered transcripts with adjusted *p*-values < 0.05 as differentially expressed, and omitted those that were not expressed in at least 25% of cells in a given annotated cell type from at least one sample.

Data manipulation and visualizations were completed in R using Seurat functions and the tidyverse package (58). The log normalized data (as described above) were used to generate violin plots and UMAP feature plots. The same data were averaged within clusters and z-scored for dot plot visualization of marker gene expression. These marker genes were chosen by performing FindMarkers on each cluster of the full dataset against all other clusters, using default values. For each cluster, the 5 genes with the highest positive fold change (avg_log2FC) were chosen from significant results (p_val_adj < 0.05) of genes that were expressed in >50% nuclei in that cluster.

Gene ontology (GO) analysis was performed with PANGEA (version 1.1 beta) (59) using the GO Hierarchy Biological Processes annotation. Genes found to be differentially expressed between CT0 and CT12 in annotated fat body cells of to>Cas9 control flies were included in gene set enrichment analysis. Genes not expressed in at least 25% of annotated fat body cells in at least one sample were excluded.

### Locomotor Activity and Feeding Rhythm Analysis

Behavioral assays were conducted on ∼7 d old male flies following entrainment to 12:12 LD cycles. For locomotor activity, flies were loaded into glass tubes containing 5% sucrose and 2% agar for analysis with the Drosophila Activity Monitoring (DAM) System (DAM2, Trikinetics Inc). DAM Monitoring was conducted for 1 week at 25°C in constant dark (DD) conditions and data were acquired every minute.

For feeding behavior, flies were mouth aspirated into individual wells of Fly Liquid-Food Interaction Counter (FLIC) monitors (Sable Systems) filled with a liquid food solution consisting of a 10% sugar solution with 45 mg/L MgCl2 for increased circuit conductance. Monitors were fitted with external reservoirs to maintain adequate food levels during monitoring. Following ∼12 hours of acclimation, feeding monitoring was conducted over 6 d in DD conditions at 25°C. Data from FLIC experiments were processed using R code developed by the Pletcher Lab (60). Feeding events were defined as times when the signal amplitude 1) exceeded the baseline readings by 5 mV for a minimum of 4 consecutive 200-ms recording periods, and 2) at some point during the event, achieved a 15 mV feeding threshold above baseline readings. Each feeding event is thus comprised of ≥ four 200-ms feeding interactions, termed “licks”.

To visualize the temporal pattern of locomotor activity and feeding over the 6 d of analysis, we created group average graphs. We first normalized individual fly locomotor activity or feeding data for each fly by dividing the value from each 30 min bin by the mean activity or feeding per 30 min across the 6d experiment. We then averaged these normalized values across all flies of a given genotype.

Circadian analysis was conducted with ClockLab software (Actimetrics). For each fly, DAM beam break and FLIC lick data were binned into 30-minute intervals, and rhythm period and power were determined over 6 consecutive days in DD using Lomb-Scargle periodogram analysis. The Lomb-Scargle “Amplitude” value at the dominant period is reported as a measure of rhythm strength (power). 3 independent experiments were run for each assay, and data from individual assays were pooled for final analysis. To account for experimental variability, Lomb-Scargle power data were normalized for each experimental replicate by dividing the power value of each fly by the mean power value of the experimental control group for that replicate. Flies that died during the course of behavioral monitoring were identified via visual inspection of activity or feeding records and excluded from analysis. For statistical comparison of locomotor activity and feeding rhythm power values across genotypes, one-way ANOVA with Dunnett’s T3 multiple comparisons test was performed with GraphPad Prism 10 software.

### Metabolomics

#### Sample Collection

We conducted metabolomics analysis on 3 genotypes of flies: iso31>per^CRISPR^ (control flies lacking GAL4), to-GAL4>per^CRISPR^ (flies lacking fat body molecular clocks), and Clk856-GAL4>per^CRISPR^ (flies lacking brain molecular clocks). We collected newly eclosed flies over a 36-hr window and aged them for 6 d in narrow polystyrene vials (at a density of 25-30 flies per vial) in 12:12 LD cycles prior to sample collection. 3 groups of flies were allowed ad libitum access to cornmeal-molasses food. A fourth, time-restricted feeding group, consisted of flies of the Clk856-GAL4>per^CRISPR^ genotype manually flipped twice daily between food-containing vials (from CT0-CT12) and agar-only vials (from CT12-CT24).

Following aging, flies were transferred to DD conditions for sample collection every 2 hours over 2 consecutive days. Each sample consisted of 30 pooled fly bodies. Flies were first CO_2_ anesthetized and collected in a 1.5 mL microcentrifuge tube for flash freezing in liquid nitrogen. They were then vortexed at high speed and passed through an ice-cold sieve (710 μm; Cole-Parmer EW-59984-07) to separate heads from bodies. The bodies were then transferred to a new 1.5 mL microcentrifuge tube and stored at - 80°C. We repeated this protocol on two separate occasions to generate 2 experimental replicates, each consisting of 24 samples per treatment group at 2-hr resolution over 2 days. The sampling frequency and replicate number were chosen to optimize detection of circadianly-regulated metabolites (61).

#### Gas Chromatography-Time of Flight Mass Spectrometry GC-TOF MS

Collected samples were sent to the West Coast Metabolomics Center (Davis, CA) for primary metabolite GC-TOF MS analysis as previously detailed (62). Samples were extracted using 1 mL of 3:3:2 ACN:IPA:H_2_O (v/v/v). Half of the sample was dried to completeness and then derivatized using 10 μl of 40 mg/mL of methoxyamine in pyridine. They were then shaken at 30°C for 1.5 hours. Then 91 μl of a N-Methyl-N-(trimethylsilyl)trifluoroacetamide with Fatty Acid Methyl Ester (MSTFA + FAME) mixture was added to each sample and they were shaken at 37°C for 0.5 hours to finish derivatization. Derivatized samples were vialed, capped, and injected into an Agilent 7890A gas chromatograph (GC) coupled with a time-of-flight mass spectrometer (LECO TOF). 0.5 μl of derivatized sample was injected using a splitless method onto a RESTEK RTX-5SIL MS column with an Intergra-Guard at 275°C with a helium flow of 1 mL/min. The GC oven was set to hold at 50°C for 1 min, then ramp up by 20°C /min to 330°C, which was held for 5 min. The transferline and EI ion source were set to 280°C and 250°C, respectively. Mass spectrometer parameters were set to collect data from 85 m/z to 500 m/z at an acquisition rate of 17 spectra/sec.

#### Data Processing and Metabolite Identification

ChromaTOF vs. 2.32 was used for data preprocessing without smoothing, 3 s peak width, baseline subtraction just above the noise level, and automatic mass spectral deconvolution and peak detection at signal/noise levels of 5:1 throughout the chromatogram. Apex masses were reported for use with the BinBase algorithm. Resulting *.txt files were exported to a data server with absolute spectra intensities and further processed by a filtering algorithm implemented in the metabolomics BinBase database. The BinBase algorithm (rtx5) used the settings: validity of chromatogram (<10 peaks with intensity>10^7 counts per s), unbiased retention index marker detection (MS similarity>800, validity of intensity range for high m/z marker ions), retention index calculation by 5^th^ order polynomial regression. Spectra were cut to 5% base peak abundance and matched to database entries from most to least abundant spectra using the following matching filters: retention index window ± 2,000 units (equivalent to about ± 2 s retention time), validation of unique ions and apex masses (unique ion must be included in apexing masses and present at >3% of base peak abundance), mass spectrum similarity must fit criteria dependent on peak purity and signal/noise ratios and a final isomer filter. Features found in less than 10% of experimental samples or with peak intensity less than 3-fold of that found in blank samples were removed from analysis. Systematic error reduction by denoising autoencoder (SERDA) normalization was performed on raw data to account for technical variation (63).

#### Metabolomics Data Analysis

To control for differences across replicates, SERDA-normalized metabolite abundance values were median-fold change adjusted across timepoints within each replicate. The Bioconductor package Rhythmicity Analysis Incorporating Nonparametric methods (RAIN v1.44.0) (39) was then used on median-fold change values to identify metabolites with rhythmic abundance rhythms at a period of 24 hours and to determine the phase of peak abundance. Individual metabolite *p*-values are automatically corrected in RAIN for the multiple waveforms and phases tested with the Benjamini-Hochberg method (64). We further corrected for the multiple metabolites tested within each group using the R p.adjust function with the Benjamini-Hochberg method to produce q-values.

## Acknowledgements

This work was supported by the Division of Integrative Organismal Systems of the National Science Foundation, CAREER Award 1942167 to D.J.C. We thank Dr. Brigitte Dauwalder, Dr. Amita Sehgal, Dr. Mimi Shirasu-Hiza, and the Bloomington Drosophila Stock Center (NIH P40OD018537) for fly stocks, and Dr. Amita Sehgal for antibodies.

