## Supplementary figures and images for "Circadian Rhythms of Whole-Body Metabolite Abundance in *Drosophila* are Largely Driven by Time of Feeding"

### Supplemental Figure 1

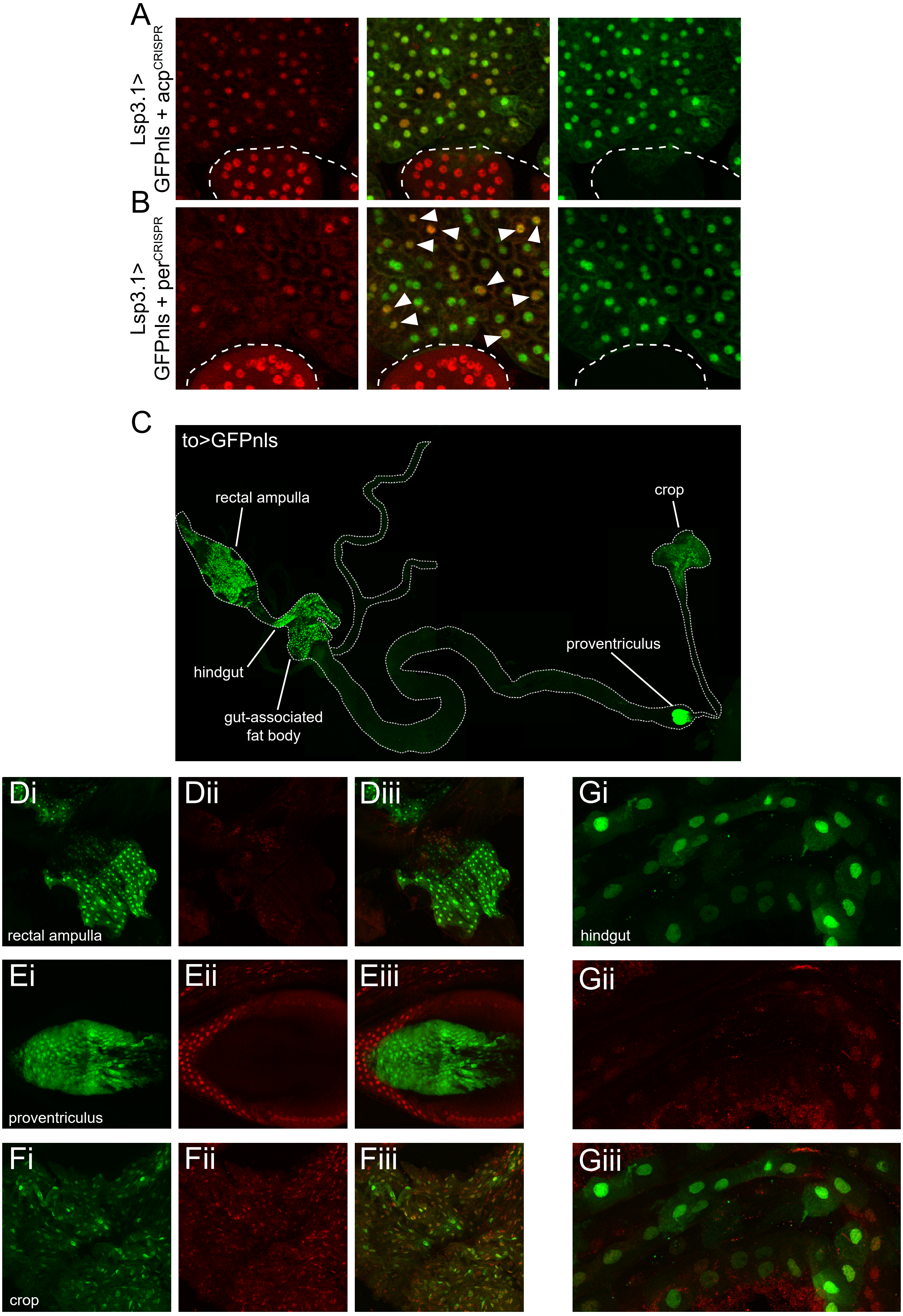

### Supplemental Figure 2

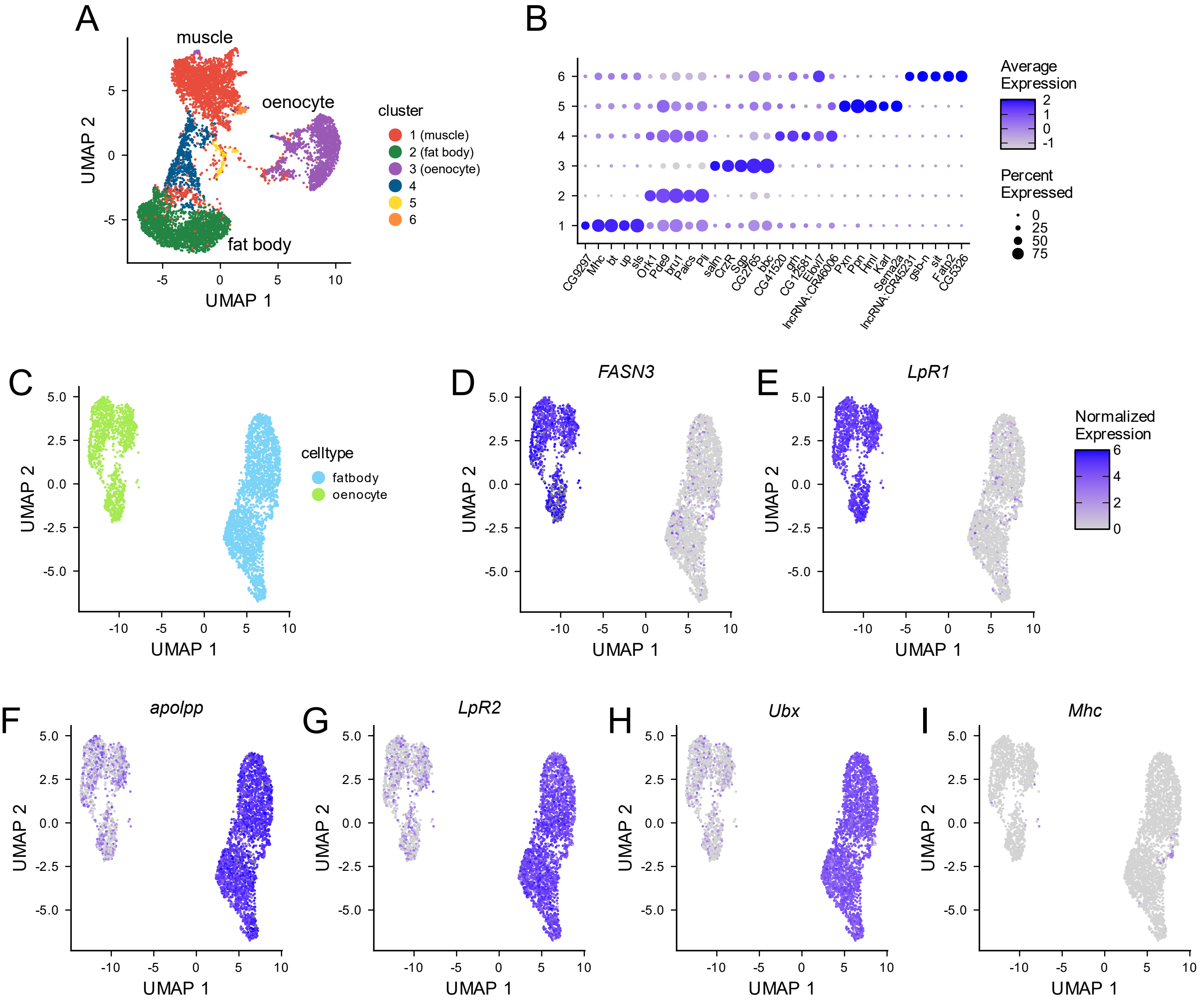
